# Neural support of manual preference revealed by BOLD variations during right and left finger-tapping in a sample of 287 healthy adults balanced for handedness

**DOI:** 10.1101/2020.09.15.297499

**Authors:** Nathalie Tzourio-Mazoyer, Loïc Labache, Laure Zago, Isabelle Hesling, Bernard Mazoyer

## Abstract

We have identified the brain areas involved in Manual Preference (MP) in 143 left-handers (LH) and 144 right-handers (RH)). First, we selected the pairs of homotopic regions of interest (hROIs) of the AICHA atlas with significant contralateral activation and asymmetry during the right-hand and the left-hand Finger-Tapping (FT) both in RH and LH. Thirteen hROIs were selected, including the primary and secondary sensorimotor, and premotor cortices, thalamus, dorsal putamen and cerebellar lobule IV. Both contralateral activations and ipsilateral deactivations (reversed for the cerebellum) were seen in primary motor and somatosensory areas, with stronger asymmetries when the preferred hand was used. Comparing the prediction of MP with different combinations of BOLD variations in these 13 hROIs, the differences between movement of the preferred hand versus that of the non-preferred hand within the contralateral and/or ipsilateral cortices of 11 hROIS performed best at explaining handedness distribution, Handedness is thus supported by: 1-between-hand variations of ipsilateral deactivations of hand primary sensorimotor and secondary somatosensory cortices and 2-variations in regions showing the same profile in left and right-handers during the right or left FT. The present study demonstrates that right and left-handedness are not based on mirrored organization of hand control areas.

## Introduction

Although handedness is a trait that has been extensively studied in humans from a variety of perspectives, whether genetic, behavioral, or sociological, it has been recently pointed out that its neural support is still to be established (Andersen & Siebner, 2018). The same observation was made by Mac Manus who recently wrote: *“Surprisingly, the nature of handedness itself has been little looked at using fMRI, the self-evident difference between the two hands being studied surprisingly rarely, despite a general recognition that left-handers are less lateralized than are right-handers”* (McManus, 2019). This raises the issue of what could possibly impede brain fMRI studies of handedness considering that there are no practical limitations in performing hand movement in the magnet.

Truly, the low prevalence of left-handedness in the population (close to 10%, (Papadatou-Pastou et al., 2019)) makes it difficult to recruit large samples of left-handed individuals. But another major difficulty in assessing the neural support of manual preference, as compared to other lateralized functions such as language, is that asymmetries measured during hand motor tasks are the direct reflection of the asymmetrical neuroanatomical organization of the motor system. Actually, 90% of white matter fibres conducting messages from the motor cortex to the medulla through the pyramidal tract are crossing at the level of the protuberance, leading to a strongly asymmetrical activity (Lemon, 2008). Therefore, asymmetries during hand movement tasks are strongly conditioned by neuroanatomy, a situation which is very different from that of language tasks that are underpinned by areas that do not have such a pattern of asymmetrical neuroanatomical organization and whose asymmetries of activation are markers of hemispheric dominance (Dym, Burns, Freeman, & Lipton, 2011). In other words, during hand movement tasks, the asymmetry favoring the hemisphere contralateral to the moving hand is not a direct measure of hemispheric dominance which may explain why no evidence for differences in asymmetry or contralateral activations between Left-Handers (LH) and Right-Handers (RH) has been revealed so far.

Taking a different perspective, Hayashi et al. (2008) have targeted the strength of deactivation in the primary motor cortex ipsilateral to the hand movement as a possible support of manual preference, grounding their hypothesis on electrophysiological studies showing that the ipsilateral activity integrates both the local activity and the transcallosal influences coming from homotopic contralateral areas (Hayashi et al., 2008). In their work, the authors demonstrated that, in RH, the ipsilateral deactivation of the motor cortex strengthens with increasing movement frequency, which led them to conclude that *“the dominance of the left primary motor cortex during right hand movement is both ipsilateral innervation and transcallosal inhibition in right-handed individuals”*.

In a previous work (Tzourio-Mazoyer et al., 2015), we pursued this line of thinking by questioning how the ipsilateral and contralateral primary hand motor cortex activity during Finger Tapping Task (FTT) varied with handedness. In agreement with Hayashi’s hypothesis, we did not find any difference between LH and RH regarding their contralateral activation, whichever the hand in action. Interestingly, we observed a difference in ipsilateral deactivation of the primary motor cortex between the movement of the preferred hand and that of the non-preferred hand, supporting again Hayashi’s hypothesis (Tzourio-Mazoyer et al., 2015). In addition, looking at another marker of handedness, namely manual ability asymmetry, we observed that the stronger the manual ability, the larger the difference in ipsilateral primary motor cortex deactivation when comparing the preferred hand movement to that of the non-preferred hand. In other words, it was the difference in the modulation of ipsilateral activity by the moving hand that was a marker of manual preference rather than a difference in contralateral minus ipsilateral asymmetries between the preferred and non-preferred hand movement.

However, the difference in ipsilateral deactivation in the primary motor cortex only explained a small part of the variance of the asymmetry of manual ability (Tzourio-Mazoyer et al., 2015) and we think that this may be due to the fact that investigations were limited to the sole primary motor cortex. Actually, the somatosensory cortex could also be an actor of the neural underpinnings of manual preference. For example, Hluschluck et al. (2006) demonstrated with magnetoencephalography (MEG) that the primary sensory cortex was characterized by a decreased ipsilateral activity in relation with the sensory feedback, and they proposed that this inhibition of the primary sensory cortex results from transcallosal inhibition that, in turn, is responsible for primary motor cortex deactivation (Hlushchuk & Hari, 2006).

In the present work, we thus extended the investigation of regional activity differences between the preferred and the non-preferred hand movements to the set of cortical areas involved in the hand sensory-motor control. Actually, as clearly demonstrated by the meta-analysis conducted by Witt et al. (2008) regarding the neural bases of FTT, the areas involved in this simple motor task constitute a large-scale network (Witt, Laird, & Meyerand, 2008).

To complete this investigation of the handedness neural support, we analysed a sample, balanced for handedness and sex, of 287 individuals of the BIL&GIN database (specifically acquired to investigate the neural support of Hemispheric Specialization (Mazoyer et al., 2016)), who completed both a Right Finger Tapping Task (RFT) and a Left Finger Tapping Task (LFT). In order to identify areas whose activity were dependent on the moving hand, we selected the cortical and the subcortical regions showing activation and asymmetry favouring the hemisphere contralateral to the moving hand during both the LFT and RFT, in RH and LH. We also included the cerebellar regions that were mapped in all individuals and which showed ipsilateral activation and asymmetry, during both the LFT and RFT, in RH and LH. Having identified a restricted set of hROIs this way, we then described their regional activity and asymmetry during right and left hand movement in RH and in LH. Second, we questioned whether, in these regions, the contralateral and ipsilateral BOLD variations between the preferred and the non-preferred hand movement or the variations in asymmetries with the moving hand, explained the distribution of handedness, and finally searched for regions which were part of the best explanatory model of this distribution.

## Material and methods

### Participants

The Basse-Normandie Ethic Committee approved the study protocol. All participants gave their informed written consent and received an allowance for their participation. All participants were free of brain abnormality as assessed by an inspection of their structural T1-MRI scans by a trained radiologist. We selected a group of 287 healthy participants from the BIL&GIN database (Mazoyer et al., 2016), including 142 women and 143 LH (69 women), who self-reported their manual preference (MP) as being right-hand or left hand. None of these individuals declared him·her·self as being a forced right-hander. Among these 287 participants, 284 were included in the previous analysis of primary hand motor area activity (M1) (Tzourio-Mazoyer et al., 2015).

The sample mean age was 26 years (SD = 6 years). LH were two years younger than RH (RH 26.8 ± 6 y; LH: 24.4 ± 6, p = 0.0007), women being one year younger than men (women 24.9 ± 5; men: 26.2 ± 7, p = 0.076) without gender x MP interaction (p = 0.50).

Participants reporting themselves as RH had an asymmetry of manual ability asymmetry measured with the finger tapping task in a limited time ((Peters & Durding, 1978) ([(right number of taps - left number of taps) / (left + right number of taps)] x 100)) of 6.26 (SD = 4.3), and their mean Edinburgh score was 93 (SD = 11). Participants reporting themselves as LH had a mean normalized finger tapping test asymmetry of −2.62 (SD = 3.9), and their mean Edinburgh score was −66 (SD = 38).

### Image acquisition

#### Structural imaging

Structural images were acquired using the same 3T Philips Intera Achieva MRI scanner including a high-resolution T1-weighted volume (T1w, sequence parameters: TR, 20 ms; TE, 4.6 ms; flip angle, 10°; inversion time, 800 ms; turbo field echo factor, 65; sense factor, 2; field of view, 256 × 256 × 180 mm^3^; isotropic voxel size 1 × 1 × 1 mm^3^). T2* -weighted multi-slice images were also acquired (T2*-FFE, fast field echo sequence parameters: TR = 3,500 ms; TE = 35 ms; flip angle = 90°; sense factor = 2; 70 axial slices; 2 x 2 x 2 mm^3^ isotropic voxel size).

#### Functional images acquisition

The task-related fMRI paradigm randomly alternated six 12-s blocks of finger tapping (3 Right Finger Tapping (RFT) and 3 Left Right Finger Tapping (LFT)) with six 12-s blocks of a central fixation crosshair reference task within a run that also included 4 blocks of 16-s visually guided saccadic eye movements (VGS) along with 4 blocks of 16-s reference central fixation task. Functional images were acquired with a T2*-weighted echo-planar sequence (T2*-EPI; 72 volumes; TR = 2 s; TE = 35 ms; flip angle = 80°; 31 axial slices; 3.75 mm^3^ isotropic voxel size) covering the same field of view as the T2*-FFE acquisition.

During the finger tapping tasks, the participant held a fiber optic response pad in each hand (Current Designs Inc, Philadelphia, PA, USA). Depending on the orientation of a symbolic cue presented at the center of the screen (arrowhead ‘>‘ or ‘<‘), the participant had to tap his right or left index finger on the response pad at 2.0 Hz as regularly as possible. Participants were instructed to perform this rhythmic finger-tapping task as long as the visual cue (> or <) was displayed (i.e., 12 s). The finger-tapping task was alternated with a reference task where the participants had to fixate a central crosshair. Both arrowheads and crosshair covered the same 0.8° x 0.8° visual angle. Motor responses from all but one participant were collected from either hand using the two fiber optic response pads. Before scanning, participants were trained to perform the finger-tapping tasks with the help of a metronome set at a frequency of 2 Hz.

### Image analysis

#### Functional imaging analysis

For each participant, (1) the T2*-FFE volume was rigidly registered to the T1-MRI; (2) the T1w volume was segmented into three brain tissue classes (grey matter, white matter, and cerebrospinal fluid; and (3) the T1-MRI scans were normalized to the BIL&GIN template including 301 volunteers from the BIL&GIN database (aligned to the MNI space) using the SPM12 “normalise” procedure with otherwise default parameters.

First, a 6-mm full width at half maximum (Gaussian filter) was applied to each run volume. Global linear modelling (statistical parametric mapping (SPM), http://www.fil.ion.ucl.ac.uk/spm/) was used for processing the task-related fMRI data. Next, three regressors were included in a general linear model. The right finger tapping task (RFT) regressor included the 3 blocks of right hand finger tapping and their 3 reference blocks, the left finger tapping task (LFT) regressor included the 3 blocks of the left hand finger tapping and their 3 reference blocks obtained during cross fixation, the saccadic eye movement task regressor included the 4 blocks of VGS and their 4 reference blocks.

Each regressor was constructed with the blocks of interest modeled by boxcar functions corresponding to paradigm timing and convolved with a standard hemodynamic response function. The multiple regression method allowed for obtaining estimates of activity levels for each task (RFT, LFT, VGS). In the present work we only consider LFT and RFT.

BOLD signal variations of RFT and LFT were measured in 185 pairs of functionally defined hROIs of the AICHA atlas (Joliot et al., 2015) adapted to SPM12 (excluding 7 hROI pairs belonging to the orbital and inferior-temporal parts of the brain in which signals were reduced due to susceptibility artefacts) by averaging the values of all voxels located within each hROI volume. Note that AICHA atlas was selected because it provides pairs of regions that are functionally homotopic and thus well suited to measure functional asymmetries.

In addition to the hROIs of the AICHA atlas, we included two cerebellar ROIs extracted from the Schmahmann atlas of the cerebellum located at the upper part of the cerebellum that were included in the mask of voxels common to all individuals of the sample in the stereotaxic space. Labelled cerebellum Cer4-5 and Cer3, these ROIs are located in the culmen of the cerebellum, on the upper and internal part of the cerebellar hemispheres (Schmahmann et al., 1999).

### Statistical analysis

Unless otherwise specified, statistical analyses were performed using the JMP Pro15 software package (www.jmp.com, SAS Institute Inc., 2018).

#### Behavioural control of right and left FTT execution during fMRI

Using a repeated-measures analysis of variance (ANOVA), we checked that the actual tapping frequency during fMRI acquisition did not differ between RFT and LFT. We also checked possible effects of age, educational level and sex, as well as interactions between the side of FTT with sex or manual preference (MP).

##### Identification of hROIs showing activation contralateral to the moving hand and having a significant asymmetry, both in RH and in LH

We conducted a conjunction analysis to select among the 185 hROIs of the AICHA atlas plus the 2 cerebellar ROIs (included in the mask common to the 287 participants) the hROIs exhibiting BOLD signal variations that were: 1-significantly positive and significantly larger on the left than on their right counterparts during RFT (larger on the right for the cerebellar hROIs); 2-significantly positive and significantly larger on the right than on their left counterparts during the LFT (larger on the left for the cerebellar ROIs). An hROI was thus selected whenever it was significantly activated and asymmetrical in a given contrast using a significance threshold set to p < 0.05. Accordingly, the significance threshold for the conjunction of activation and asymmetry in a given task was 0.05 x 0.05 = 2.5 x 10^−3^, and when considering the two tasks, the overall significance threshold for the conjunction of the conjunction analyses was thus 6.25 x 10^−6^ = (2.5 x 10^−3^)^2^. We carried out this selection in RH and LH separately. Finally, the selected hROIs were those common to RH and LH, leading the final significance threshold to p = 3.9 x 10^−11^ = (6.25 x 10-^6^)^2^.

##### Profiles of activity and asymmetries in the so-defined set of hROIs, and effect of movement side in RH and LH

In RH, we first described the anatomical location and FTT-induced BOLD variations of the selected hROIs in the hemisphere contralateral and in the hemisphere ipsilateral to the right (preferred) hand movement and their asymmetry.

We then investigated, only in RH, the variations of hROIs activations and asymmetries profiles depending on the moving hand. In order to do so, we completed a MANOVA with repeated measures on the 13 hROIs with a Task main effect, defined as moving the right hand or the left hand, and their interaction (Task by hROI).

The same analysis was completed in LH.

#### Prediction of manual preference from regional BOLD variations during hand movements: comparison of 4 models

We compared 4 models as regards their capacity at explaining and predicting manual preference, each model having its specific set of 26 explanatory variables obtained by different combinations of the BOLD variations during each hand movement in the 13 cortical, subcortical and cerebellar ROIs in each hemisphere.

Model I was based on our working hypothesis stipulating that the neural underpinning of handedness is a modulation of contralateral and ipsilateral BOLD variations that depends on the moving **hand laterality** (i.e. preferred or non-preferred hand);

Model II also examined both contralateral and ipsilateral variations as model I, but considered that it is the moving **hand side** (i.e. left or right) that modulates these variations rather than hand laterality;

Model III and model IV considered regional asymmetries values (contralateral minus ipsilateral) measured during both hands movements, one looking at the moving **hand side** (model III) the other at the moving **hand laterality** (model IV).

These models can be formalized as follows:

- Model I: [Contralateral(preferredFTT - non-preferredFTT)], and [Ipsilateral(preferredF-TT - non-preferredFTT)]
- Model II: Model II : [Contralateral(RFT - LFT)], and [Ipsilateral(RFT - LFT)]
- Model III: Model III: [RFT(Contralateral - Ipsilateral)], and [LFT(Contralateral - Ipsilateral)]
- Model IV: [PreferredFTT(Contralateral - Ipsilateral)], and [non-preferredFTT(Contralateral - Ipsilateral)]

Each of these 4 models was first optimized in terms of which variables were to be considered as significantly contributing to handedness prediction using a descending stepwise logistic regression with handedness as the dependent variable and the Akaike’s Information Criterion (AIC) as a stopping rule. Goodness of fit of each model was measured using both the R2 and adjusted-R2 values, and 95% confidence intervals (95%CI) were computed for both statistics using the formula provided by Cohen et al. pp 179-181 (Cohen, Cohen, West, & Aiken, 2003). Chance-corrected agreement between the model-predicted MP and the actual MP was then measured using the Kappa statistic, and the 95%CI of the Kappa value was also computed. Models were then compared for each of the 3 statistics (R^2^, adjusted-R^2^, and Kappa) based on their respective 95%CI. For the best identified model, we then described how each explanatory variable contributed to handedness prediction.

## Results

### Behavioural control of FTT execution during fMRI

Neither movement side, MP, sex or their interaction had significant effects on the mean frequencies of finger tapping recorded during fMRI acquisition (RH: RFT= 2.17 ± 0.38 Hz, LFT= 2.17 ± 0.39 Hz, LH : RFT= 2.24 ± 0.43 Hz, LFT= 2.26 ± 0.41 Hz). Note also that there was no effect of age on these frequencies.

### Identification of hROIs showing activation contralateral to the moving hand and having a significant asymmetry, in both RH and LH

The conjunction analysis uncovered 12 supratentorial and one cerebellar hROIs showing a BOLD activity shifting side with the moving hand. The 12 supratentorial regions were significantly activated contralaterally to the moving hand and showed a shift in asymmetry when switching the moving hand in both groups (Figure 1), while the cerebellar hROI exhibited the reverse pattern.

#### Anatomical localization of the 13 hROIs

The coordinates of the center of mass of each selected hROI are provided in Table 3.

**Figure 1.**
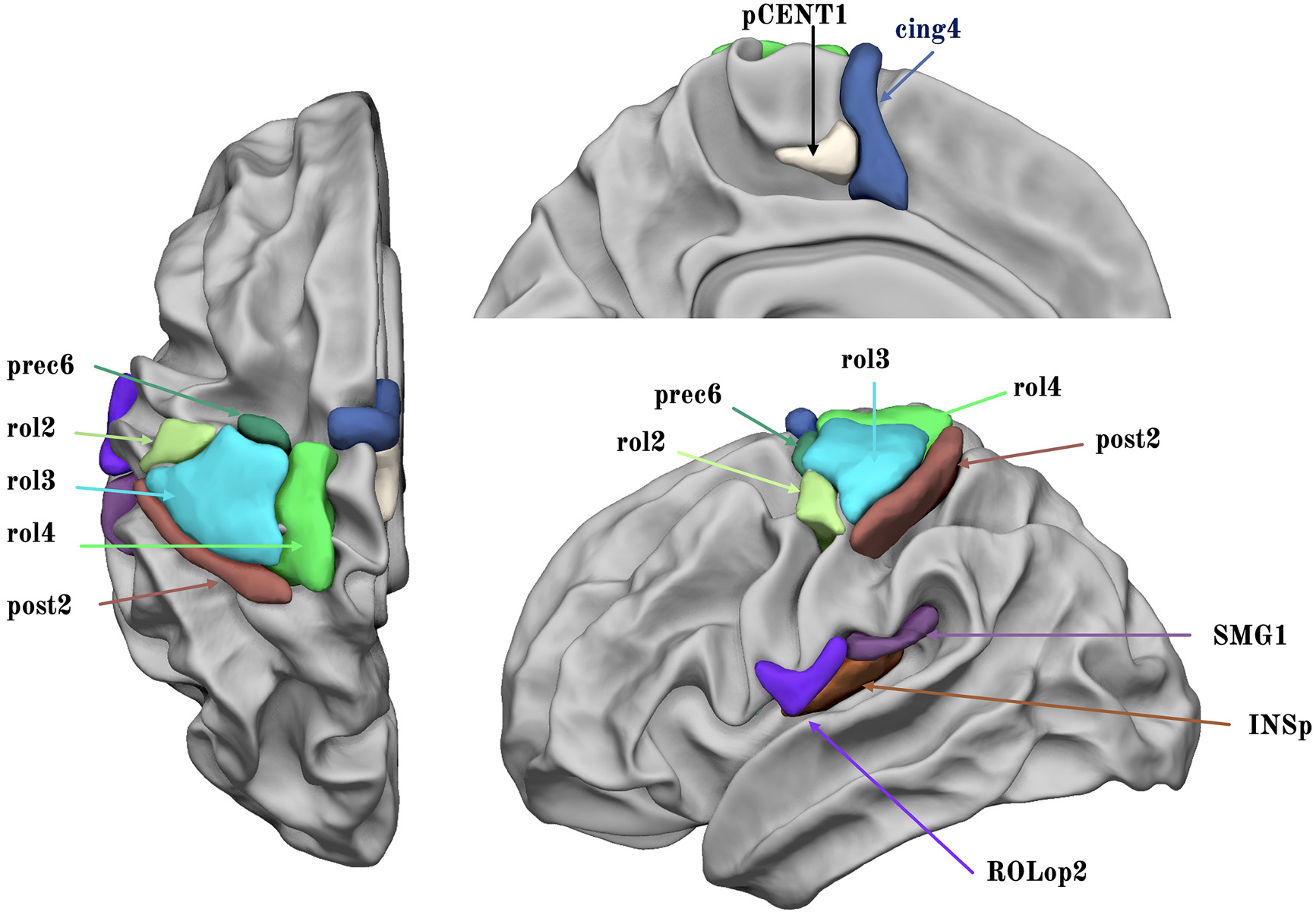
Display of the 12 supratentorial hROIs contralaterally activated and having significantly larger contralateral minus ipsilateral asymmetry in both right and left finger tapping tasks. (Abbreviations: Rolandic sulcus: rol, cingulate sulcus: cing, precentral sulcus: prec, paracentral gyrus: pCENT1, Rolandic operculum: ROLop, supramarginal gyrus: SMG, posterior insula: INSp1).

At the cortical level, the upper Rolandic sulcus areas included 3 hROIs: rol4, rol3, rol2. The hROI rol3 was located precisely at the hand rolandic genu corresponding to the location of the hand motor area, while rol2 was located immediately ventrally and rol4 immediately dorsally to rol3.

A premotor region was located in the precentral sulcus (prec6) immediately anterior to rol3.

Medially, the hROIs corresponding to SMA proper and to the pre-SMA were part of the set of hand motor response areas (pCENT1, cing4).

The hand primary somatosensory cortex along the postcentral sulcus at the posterior border of rol3, corresponded to the hROI post2 overlapping area 3b (Hlushchuk, Simões-Franklin, Nangini, & Hari, 2015), while the secondary somatosensory cortex corresponded to a bunch of adjacent hROIs centred by the posterior insula hROIs (INSp1) and extending to the rolandic and parietal operculum (ROLop2, SMG1).

Subcortical areas included the anterior two third of the thalamus (THA5) and the posterior part of the putamen (PUT3). As upper mentioned, the cerebellar hROI was located in the IVth lobule (see Figure 5).

### Patterns of regional bold variations and asymmetries during right- and left-hand movements in right-handers. Effects and interaction of hand movement side and hROI

Contralateral and ipsilateral regional BOLD variations in the 13 hROIs are shown in Figure 2. The MANOVA of contralateral BOLD variations showed significant hROI (p<0.0001) and Task (p = 0.0003) main effects and hROI by Task interaction (p<0.0001).

**Figure 2.**
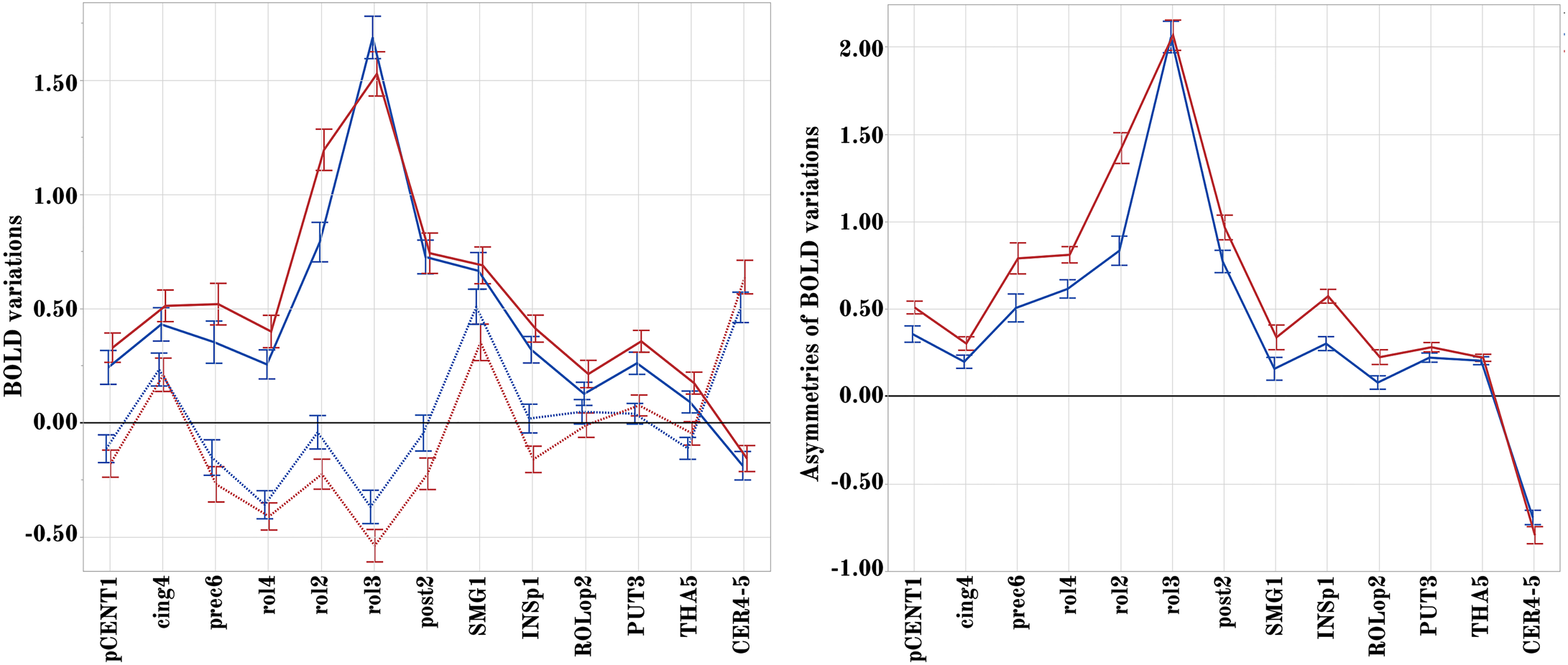
Regional BOLD signal variations and asymmetries during movements of the right (preferred) and left (non-preferred) hands in RH. Left panel: contralateral (RFT: red, LFT: blue) and ipsilateral BOLD signal variations (dashed line) in the 13h ROIs. Right panel: regional asymmetry (contralateral minus ipsilateral) of BOLD signal variations in the same hROIs (RFT: red, LFT: blue). Values are group means with error bars showing 95% confidence intervals. FT: finger tapping.

The hROI main effect corresponded to larger contralateral activations in the hand primary motor (rol3) and adjacent motor area (rol2), and somatosensory hROIs (post2), than in the other hROIs, whatever the hand moved. The Task main effect was due to larger activations on average during movement of the right hand than during movement of the left hand. The hROI x Task interaction was due to significantly larger activations in rol3 during movement of the left hand than during movement of the right hand while the reverse pattern was observed in most other hROIs.

The MANOVA of ipsilateral activity also showed significant hROI (p<0.0001) and Task (p = 0.0003) main effects and a hROI by Task interaction (p<0.0001). The hROI main effect was due to deactivations in the rol3, rol4 and prec6 hROIs as compared to activation in SMG1 and cing4. The Task main effect was due to lower values during RFT than during LFT. The hROI x Task interaction was due to significantly larger deactivations in rol2, rol3, rol4, post2, and INsp1 during movement of the (preferred) right hand than during movement of the left hand as opposed to the absence of difference in cing4 and subcortical areas.

#### Asymmetries

The MANOVA on asymmetries evidenced significant hROI (p<0.0001) and Task (p<0.0001) main effects and hROI by Task interaction (p<0.0001).

The hROI main effect was related to strong variations of the strength of asymmetries across areas, with stronger values in hand primary motor, premotor and somatosensory areas (rol3, rol2, rol4 prec6 and post2) compared to all other hROIs (see Figure 2, right). The Task main effect corresponded to stronger asymmetries during RFT than during LFT, in all hROIs but rol3 and subcortical areas (Figure 2) which was the cause of the task x hROI significant interaction. Note the large asymmetry favoring the ipsilateral side for the cerebellar ROI.

### Patterns of regional bold variations and asymmetries during right- and left-hand movements in left-handers. Effects and interaction of hand movement side and hROI

Contralateral and ipsilateral BOLD variations are shown in Figure 3.

**Figure 3.**
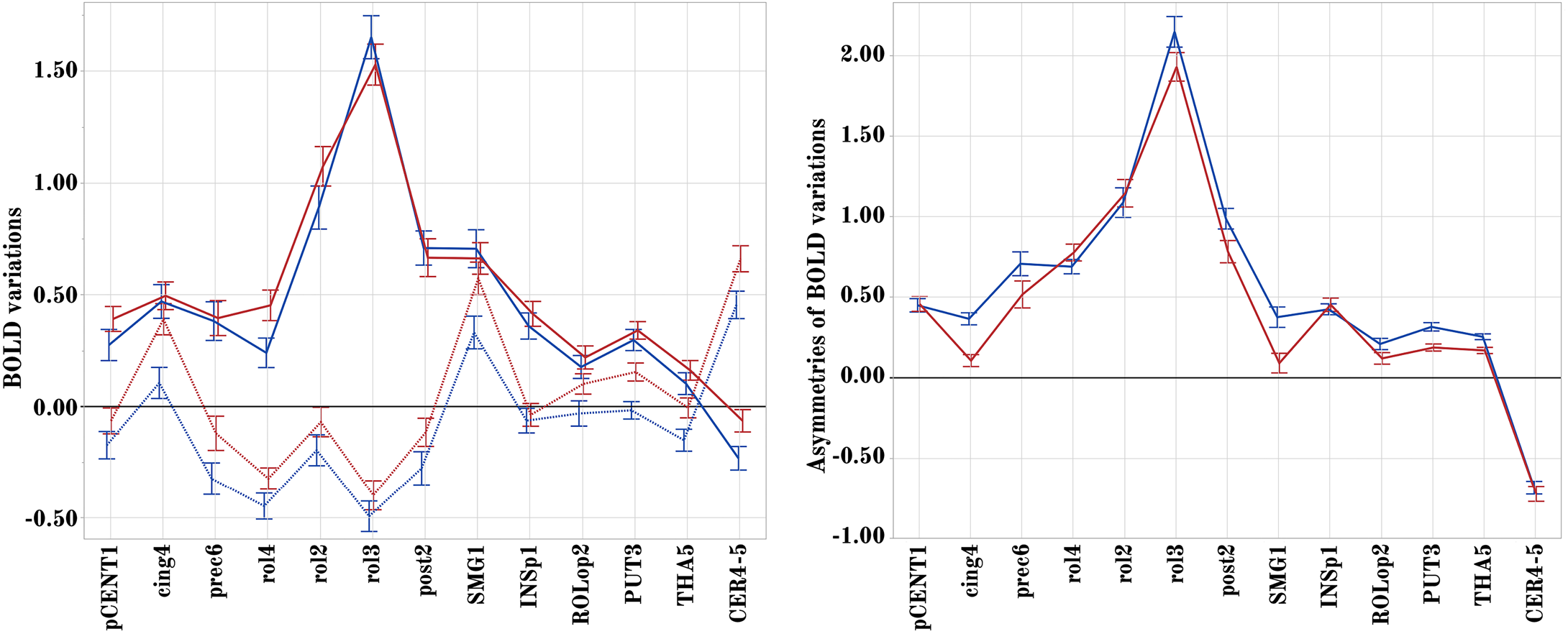
Regional BOLD signal variations during movements of the left (preferred) and right (non-preferred) hands in LH. Left panel: contralateral (RFT: red, LFT: blue) and ipsilateral BOLD signal variations in the 13h ROIs (dashed lines). Right panel: regional asymmetry (contralateral minus ipsilateral) of BOLD signal variations in the same hROIs (RFT: red, LFT: blue). Values are group means with error bars showing 95% confidence intervals. FT: finger tapping.

MANOVA of contralateral activity in LH showed significant hROI main effect (p<0.0001) and hROI by Task interaction (p<0.0001), but, contrary to RH, no significant Task main effect (p = 0.078). The absence of a significant Task effect in LH was related to comparable contralateral activations during RFT and LFT in most hROIs (see Figure 3 left panel). However, rol2, rol4 and paraCENT1 contralateral activations and CER4_5 ipsilateral activation were larger during LFT than during RFT (p-values), explaining the significant Task x hROI interaction.

MANOVA of ipsilateral activity showed significant hROI (p<0.0001) and Task (p<0.0001) main effects and hROI by Task interaction (p<0.0001). Similar to RH, LH had stronger ipsilateral deactivation during their preferred (left) hand movement, and the significant Task x hROI interaction was due to a non-significant difference between the 2 tasks regarding INSp1 variations as opposed to all other hROIs.

#### Asymmetries

As for RH, the MANOVA on asymmetries evidenced significant hROI (p<0.0001) and Task (p<0.0001) main effects and Task by hROI interaction (p<0.0001). The main Task effect was related to larger mean asymmetry during left hand movement while the Task by hROI interaction was related to an absence of difference in pCENT1, INSP1, rol2, rol4 and CER4_5 (see Figure 3 right panel). Note again the large asymmetry favoring the ipsilateral side for the cerebellar ROI.

### Explaining handedness

#### Comparison of the 4 models

Results are summarized in Table 1.

**Table 1.** Goodness of fit (R^2^ and Adjusted-R^2^) and concordance (Kappa) between model-predicted and actual manual preferences for 4 different models (see text for description of models). 95%CI: 95% confidence interval.

| Model | Nvar | R<sup>2</sup> [95%CI] | Adjusted-R<sup>2</sup> [95%CI] | Kappa [95%CI] |
| --- | --- | --- | --- | --- |
| I | 8 | 0.336 [0.250, 0.422] | 0.317 [0.229, 0.405] | 0.596 [0.503, 0.688] |
| II | 17 | 0.590 [0.522, 0.658] | 0.590 [0.492, 0.635] | 0.728 [0.649, 0.807] |
| III | 12 | 0.383 [0.299, 0.467] | 0.356 [0.268, 0.443] | 0.630 [0.540, 0.720] |
| IV | 15 | 0.811 [0.775, 0.848] | 0.801 [0.762, 0.840] | 0.881 [0.827, 0.935] |

Model I, that includes contralateral and ipsilateral regional differences between BOLD signal variations during the preferred and during the non-preferred hand movement, significantly outperformed the 3 other models in terms of both goodness-of-fit (R^2^) and handedness prediction (Kappa).

Model IV (asymmetries during FTT of the preferred and of the non-preferred hand) was significantly better than both model II (variations of contralateral and ipsilateral activity between RFT and LFT) and model III (asymmetries of RFT and LFT) in terms of goodness-of-fit, whereas (adjusted) R^2^-values for the latter two models did not significantly differ.

However, Kappa values for models II, III, and IV, were not significantly different.

#### Best model for predicting manual preference

Model I exhibited both high R^2^ and high Kappa values, with 94% of raw concordance between actual and model-predicted MP (95% in RH and 93% in LH). According to the AIC of the stepwise descending logistic regression, the optimal set of explanatory variables for handedness included differences in BOLD variations between FTT of the preferred hand and FTT of the non-preferred hand in 16 hROIs (see Table 2). Of these 16 hROIs, 8 were contralateral and 8 were rol2 ipsilateral to the movement, including 5 pairs of homotopic hROIs (rol3, cin4, post2, INSp1, SMG1).

**Table 2.** Explanatory variables with significant contribution to the logistic regression of manual preference using model I. *Note that connections between the hemispheres and the cerebellum are crossed and therefore ipsilateral cerebellum corresponds to contralateral cortical areas.

| hROI (abbreviation) | Contralateral |  |  | Ipsilateral |  |  |
| --- | --- | --- | --- | --- | --- | --- |
|  | Khi-square | Log Worth | p | Khi-square | Log Worth | p |
| S_Rolando-3 (rol3) | 108.5 | 24.69 | <0.0001 | 5.98 | 1.84 | 0.014 |
| S_Rolando-4 (rol4) | 47.59 | 11.28 | <0.0001 |  |  |  |
| S_Rolando-2 (rol2) | 32.62 | 7.95 | <0.0001 |  |  |  |
| S_Cingulate-4 (cin4) | 10.94 | 3.02 | 0.0009 | 12.29 | 3.34 | 0.0005 |
| S_Postcentral-2 (post2) | 5.28 | 1.66 | 0.021 | 5.02 | 1.60 | 0.025 |
| G_Insula-posterior-1 (INSp1) | 10.94 | 3.02 | 0.0009 | 29.48 | 7.25 | <0.0001 |
| G_Supramarginal-1 (SMG1) | 11.48 | 3.15 | 0.0007 | 4.92 | 1.57 | 0.026 |
| G_Rolandic_Oper-2 (ROLOp2) |  |  |  | 6.68 | 2.01 | 0.0097 |
| N_Putamen-3 (PUT3) | 3.18 | 1.12 | 0.07 |  |  |  |
| N_Thalamus-5 (THA5) |  |  |  | 11.67 | 3.19 | 0.00006 |
| Cerebellum (CER4_5) |  |  |  | 18.33 | 4.73 | <0.0001 |

Contralateral motor areas (rol2, rol3, rol4) were the hROIs in which differences in regional BOLD variations had a maximum significance for MP explanation, compared to secondary somatosensory regions (INSp1, SMG1, post2) where smaller significance was observed. Ipsilateral hROIs for which differences in regional BOLD variations between the FTT of the preferred hand and of the non-preferred hand that were found to significantly contribute to MP explanation included only one of the motor area (rol3), the 3 adjacent regions of the secondary sensory cortex (INSP1, SMG1, ROLop2), INSP1 having the most significant explanatory power among all, as well as the cerebellar hROI.

Figure 4 details how handedness modulates the difference in regional BOLD signal variations between the preferred hand movement and the non-preferred hand movement for each of these 16 hROIs. This modulation appears to follow 3 different patterns across the set of 16 hROIs: the first and second patterns were characterized by differences in BOLD signals between movements of the two hands that were of opposite signs in RH and in LH, whereas the third one concerned hROIs for which this difference had the same sign but different magnitudes in RH and in LH.

**Figure 4.**
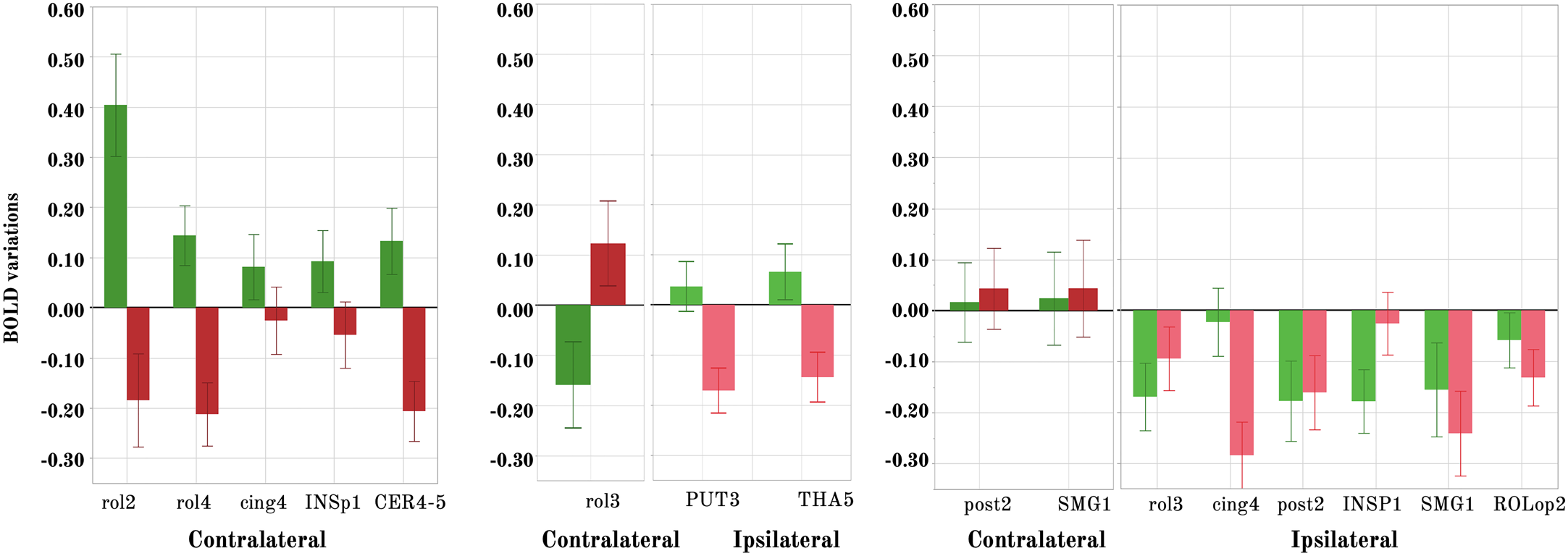
Differences between the activity during the preferred hand movement minus the non-preferred hand movement in RH (green) and LH (red) in the hROIs selected by the best model for explaining handedness distribution. Left panel: Contralateral variations in regions showing opposite effects in LH and RH, with a positive value in RH corresponding to larger activation in the left hemisphere during RFT than in the right hemisphere during LFT. Middle panel: contralateral variations showing opposite effects in LH and RH, with a positive value in rol3 for LH therefore corresponding to a larger activation in the right hemisphere during LFT than in the left hemisphere during RFT, and in PUT3 and THA5 with a stronger ipsilateral deactivation during LFT than during RFT. Right panel: Regions where a stronger contralateral activation (post2, SMG1) and a stronger ipsilateral deactivation (rol3, cing4, post2, SMG1, ROLop2) are present during the preferred hand movement. Dark colours correspond to contralateral variations and light colours to ipsilateral variations.

The first pattern (see left panel of Figure 4) concerned contralateral hROIs for which RH had larger activation during movement of their preferred hand (i.e. the right) as compared to movement of their non-preferred hand (i.e. the left), whereas LH had larger activation during movement of their non-preferred hand (i.e. the right) than during movement of their preferred hand (i.e. the left). In other words, when moving their right hand both groups showed larger contralateral activation in rol2, rol4, INSP1 and larger ipsilateral cerebellar activation in CER-4_5. In rol2 and INSP1 the larger contralateral activation during preferred hand movement compared to non-preferred hand movement was stronger in RH, while, on the opposite, the difference was stronger in LH in rol4 and ipsilateral CER4-5.

The second pattern (reverse of the first) concerned contralateral hROIs for which RH had lower activation during movement of their preferred hand compared to movement of their non-preferred hand, whereas LH had larger activation during movement of their preferred hand than during movement of their non-preferred hand (see middle panel of Figure 4).

Ipsilateral hROIs for which LH (respectively RH) showed larger (respectively lower) deactivation during movement of their preferred hand (as compared to their non-preferred hand) were considered to also exhibit this second pattern. In other terms, both groups showed larger contralateral activation in the hand motor area rol3 during their left hand movement, that was therefore of opposite signs, and rol3 contralateral variations was the most significant explanatory variable (Table 2). They also showed larger ipsilateral deactivation in subcortical areas PUT3 and THA5 when moving their left hand, the difference being larger in LH.

The third pattern concerned hROIs in which differences in BOLD variations between FFT of the preferred and of the non-preferred hand had the same sign but different magnitudes in RH and in LH (see right panel of Figure 4). For instance, FTT of the preferred hand elicited larger ipsilateral decreases than FTT of the non-preferred hand in the rol3 motor area as well as in 2 regions corresponding to the secondary somatosensory cortex (INSP1 and ROLop2), these larger ipsilateral deactivations being stronger in RH for the rol3 and INSP1 hROIs, and stronger in LH for ROLop2. In SMG1 lower ipsilateral activation was present when moving the non-preferred hand and this effect was stronger in LH. SMA, corresponding to cing4, was recruited ipsilaterally with the same magnitude for both hands in RH, but with a stronger intensity when LH moved their right hand, Finally, slightly larger contralateral activation during preferred hand movement in the hand primary sensory cortex (post2) and SMG1 were present in both groups. Although non-significant, this increase of small differences in activation in LH as compared to RH was sufficient for retaining these 2 areas when optimizing model 1.

## Discussion

The present work, that investigated in a large population balanced for handedness the neural activity associated with movement of each hand in the same individuals, led to original observations regarding the organisation of the motor control of one hand relatively to the other, and identified a set of areas which functional patterns explained 96% of the sample distribution of handedness.

Because our aim was to question how the regions involved in hand movement support handedness, we selected regions that shifted their activity asymmetry with the moving hand, thereby removing other lateralized components of the tasks. Indeed, since we measured BOLD variations during internally guided movements at a pre-learned pace in right- and left-handers it is clear that other cognitive components than motor systems are recruited to appropriately complete such a complex task. For example, the internal pacing at the learned rhythm mainly driven by the left hemisphere for overlearned rhythm (Pflug, Gompf, & Kell, 2017), rightward lateralized attentional control (Zago et al., 2017), and verbal rehearsal involving left audio-motor loop (Hesling, Labache, Joliot, & Tzourio-Mazoyer, 2019).

### A set of core hand sensorimotor areas

It is first noticeable that the set of regions activated and asymmetrical that shifted hemisphere when shifting moving hand was quite large and overlapped those of the meta-analysis conducted by Witt on finger-tapping tasks activations mapped with fMRI (Dos Santos Sequeira et al., 2006)). In this last work, gathering results of 38 different finger-tapping tasks for an ALE-based meta-analysis, the authors targeted clusters of concordance in the left hemisphere among the various experimental paradigms of finger tapping tasks involving right finger or bilateral finger movements. As shown in Figure 5A, the primary motor, premotor, secondary somatosensory, subcortical areas and cerebellar areas reported by Witt et al. were very similar to the regions identified in the present work. Interestingly, the set of regions of the present study also overlapped those disconnected by upper limb disuse and showing large pulses of spontaneous activity (Figure 5B, (Newbold et al., 2020)). We can thus consider the selection we operated as efficient to unravel the hand-movement control core brain network.

**Figure 5.**
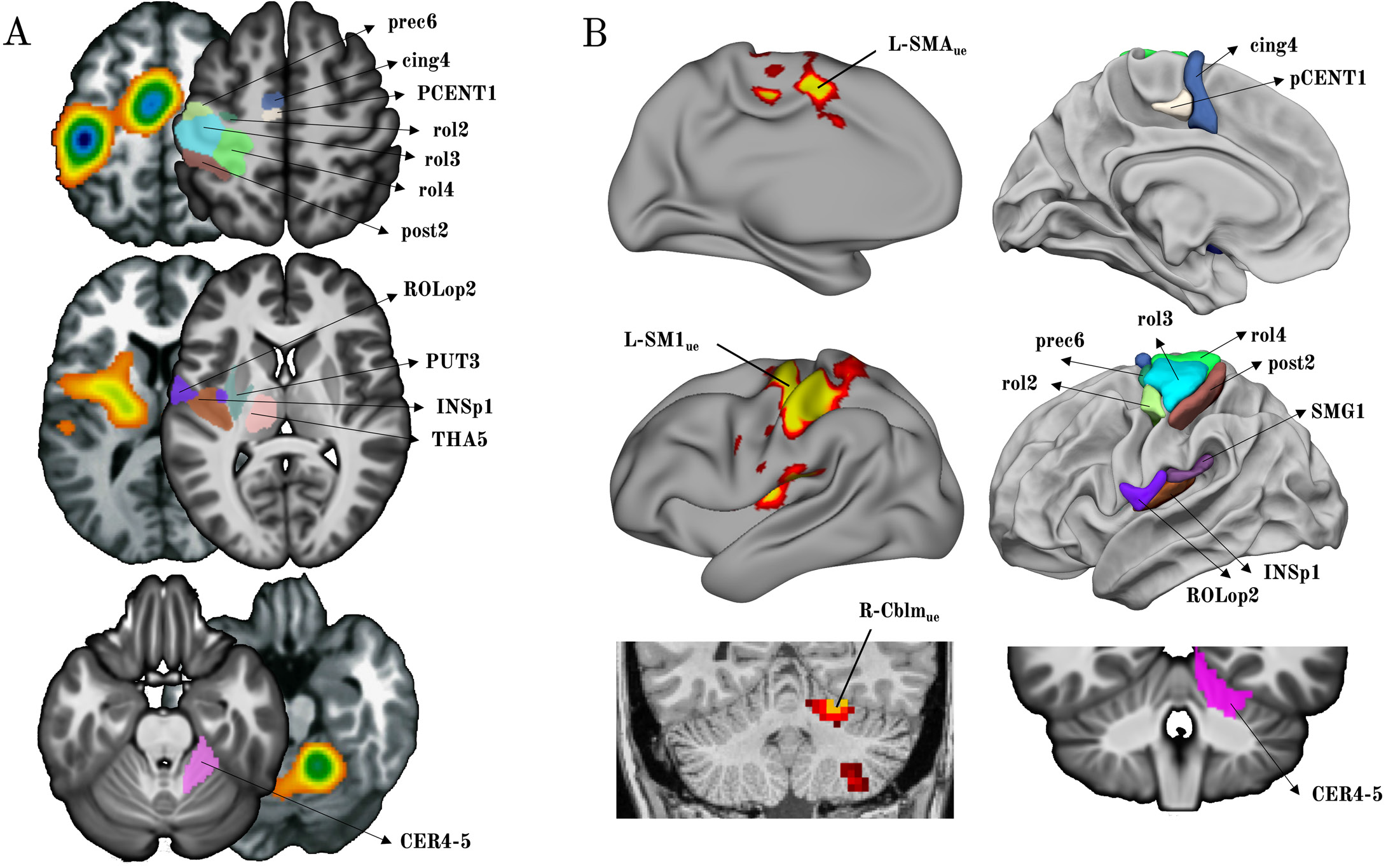
Location of the 12 hROIs showed in the left hemisphere that were selected in the present work. (A) Comparison with the map of the main effect of all finger-tapping tasks in the meta-analysis conducted by Witt (adapted from (Witt et al., 2008)). The ALE maps are shown side to side with the hROIs of the present work that are labeled on axial slices of the left hemisphere and right cerebellum (B) Comparison with the regions involved in the motor network for hand movements as explored with intrinsic connectivity by Newbold together with lateral left hemisphere presentation of the hROIs of the present work, note that in Newbold the coral slice presenting the right cerebellum is in a non-radiological convention (adapted from (Newbold et al., 2020).

More precisely, despite the coarse resolution of a large meta-analysis and the subsequent reduction of the number of independent peaks, the comparison of the MNI coordinates of the present hROIs centre of mass with those of the peaks reported in Witt’s meta-analysis allows us to make solid hypotheses on the role of the selected hROIs.

For instance, it is beyond any doubt that rol3 corresponds to the primary motor cortex considering its location at the *genu* of Rolando corresponding to M1.

As regards rol2, it has the same coordinates than the “dorsal premotor” activation reported in paced FTT tasks in Witt et al. meta-analysis ((Witt et al., 2008), see Table 3)). The cing4 hROI also has coordinates comparable with those of the region Witt et al. labeled as SMA, in accordance with its location posterior to the anterior commissure line. We also consider that the pCENT1 is part of SMA, located dorsally to cing4. As for the premotor regions coordinates of the Witt’s study, although they were not identified as separate peaks because of the low resolution of the ALE approach (the authors state: “*Rather, the clusters comprising the primary sensorimotor cortices extended anterior enough to encompass the cortical region usually defined as the dorsal premotor cortex, and the location of this extension in the left hemisphere was in agreement with the region of the left PMd identified through a previous meta-analysis study*…”), they had spatial locations in the cluster that overlaps rol2 and prec6 coordinates. The ventral part of the premotor was identified as a specific peak in Witt et al. meta-analysis that overlaps ROLop2 hROI. The post2 hROI was located precisely at the location of the hand primary sensory cortex, while not differentiated in the meta-analysis from the large sensorimotor cluster (Figure 5A). Finally, a cluster located more deeply in Witt’s study overlapped our INSp1, PUT3 and THA5 hROIs, while identical coordinates were found for the center of mass of the CER4_5 hROI and the cerebellar peak of Witt’s meta-analysis (Table3, Figure 5A).

**Table 3.** Coordinates of the center of mass of the hROIs of the present study and of the clusters identified in Witt’s meta-analysis of right finger tapping tasks (Witt et al., 2008).

| AICHA hROI | Coordinates<br>left hemisphere |  |  | Coordinates<br>*Witt et al. 2008* |  |  | Label<br>*Witt et al. 2008* |
| --- | --- | --- | --- | --- | --- | --- | --- |
|  | x | y | z | x | y | z |  |
| S_Rolando-3 (rol3) | -38.8 | -23.1 | 61.4 | -38 | -26 | 50 | M1 |
| S_Rolando-4 (rol4) | -23.1 | -28.9 | 65.3 |  |  |  |  |
| S_Rolando-2 (rol2) | -43.6 | -13.7 | 50.6 | 38 | -10 | 54 | Dorsal premotor* |
| S_Precentral-6 (prec6) | -29.8 | -11.2 | 64.7 |  |  |  |  |
| S_Cingulate-4 (cin4) | -7.8 | -6.3 | 57.4 | -4 | -8 | 52 | SMA |
| G_Paracentral-1 (pCENT1) | -6.9 | -16.8 | 50.9 |  |  |  |  |
| S_Postcentral-2 (post2) | -40.8 | -33.5 | 54.7 |  |  |  |  |
| G_Insula-posterior-1 (INSp1) | -42.1 | -19.1 | 13.7 |  |  |  |  |
| G_Supramarginal-1 (SMG1) | -54.5 | -29.5 | 21.4 | -50 | -26 | 20 | Inferior parietal |
| G_Rolandic_Oper-2 (ROlop2) | -50.6 | -9.0 | 13.9 | -52 | -2 | 10 | Ventral premotor |
| N_Putamen-3 (PUT3) | -28.0 | -6.3 | 1.8 | -22 | -8 | 4 | Basal ganglia |
| N_Thalamus-5 (THA5) | -12.4 | -19.0 | 7.0 |  |  |  |  |
| Cerebellum (CER4_5) | 14 | -49 | -18.0 | 16 | -50 | -20 |  |

### Ipsilateral inhibition is not limited to hand sensorimotor cortex

In this set of 13 hROIs the highest activations and asymmetries were found in the primary motor and primary somatosensory cortices in accordance with their pauci-synaptic connection with the sensorimotor effectors and receptors, followed, in terms of activation strength, by premotor and secondary somatosensory cortices. Asymmetries strength was related to contralateral activations but also in part to the strength of ipsilateral inhibition in homotopic regions leading to deactivations not limited to primary motor regions or to the primary sensory hand area, as it has been previously described (Hayashi et al., 2008; Hlushchuk & Hari, 2006). Notably, deactivations were observed also at distance from the ipsilateral primary motor cortex, at the location of the secondary somatosensory cortex (INSp1). The present results thus demonstrate that inhibition from the contralateral cortex towards the ipsilateral homotopic cortex through the corpus callosum resulting in ipsilateral deactivations is not limited to primary hand motor and sensory areas. Actually it can be considered that not only deactivations, but also weaker activations observed during the preferred hand than during the non-preferred hand movement, correspond to the same phenomenon present in almost all supratentorial areas in both RH and LH (see Figures 2 and 3). Inter-hemispheric inhibition from the contralateral hemisphere towards the ipsilateral hemisphere thus seems to be a global mechanism at play during single hand movement, acting either via direct callosal connections (like in primary motor regions), or spreading at distance across regions belonging to the same network as proposed by Hlushchuk et al. (Hlushchuk & Hari, 2006). Observations of split-brain patients showing long-lasting difficulties of coordination of bilateral complex hand movements after surgery (Zaidel & Sperry, 1977), support the hypothesis that interhemispheric inhibition is essential to learn or execute complex movements requiring inter-manual coordination.

### Within the set of regions involved in hand motor control, contralateral functional variations code for hand side, while ipsilateral variations code for hand laterality

It must be underlined that we observed variations in regional activity that were associated with either the moving hand side (right or left) or the moving hand laterality (preferred or non-preferred), that appeared to act upon these variations as a modulator, whatever handedness.

When moving their left hand, all individuals, whether RH or LH, had stronger contralateral activity of their contralateral (right) hand motor area, rol3, whereas when moving their right hand, it was the surrounding rol2 and rol4 contralateral activity that exhibited stronger activity. These variations of activity in ROIs centered by the hand primary motor areas and strongly connected by the corpus callosum across hemispheres, point to a spatial functional organization of the hand motor cortex that appears to have been previously overlooked. Such a coding of hand laterality in contiguous parts of the Rolandic cortex is likely to play an important role in complex bi-manual activities, an hypothesis that calls for further investigations.

On the opposite, modulation of variations of activity with hand preference mainly involved cortices ipsilateral to the moving hand, underlining the importance of interhemispheric inhibition in the setting of hand preference. This phenomenon concerned the ipsilateral hand motor area rol3, in agreement with previous works showing larger deactivation in this area during the preferred hand movement than during the non-preferred hand movement (Tzourio-Mazoyer et al., 2015; Hayashi et al., 2008). But here, we report for the first time that this phenomenon also concerned all ipsilateral hROIs encompassing the secondary somatosensory cortex, that may be due to a cascade of inhibition coming from the primary sensory cortex with which secondary sensory areas are strongly connected. Actually as shown by Hlushchuk et al. (2006), the sensory feedback results in decrease in activity due to transcallosal inhibition in primary sensory cortex that in turn results in decreased activity in primary motor areas, we propose that the same phenomenon is at play for ipsilateral secondary sensory areas. The regional differences observed here between RH and LH with a stronger increase in ipsilateral deactivations during the preferred hand movement in INSP1 for RH, and in ROLop2 in LH remain to be understood, calling for further explorations.

### Neural support of handedness

The present results show that handedness is characterized by the fact that some regions within the hand sensorimotor network have an activity profile during hand movement that depends on whether it is the preferred or the non-preferred hand that is moving, while some other areas do not experience such a modulation of their activity. These two types of variations in hROIs behaviour explain why the model we proposed, based on differences between preferred and non-preferred BOLD variations in contralateral and ipsilateral cortices, was able to explain handedness distribution to a large extent. As we have shown, other models based on hand side did not include the regions where differences related to the use of preferred and non-preferred hand were observed, while models based on asymmetry did not make it possible to differentiate between contralateral and ipsilateral variations. Actually, differences between right and left hand movements asymmetries explained only a small fraction of handedness distribution, confirming that contralateral minus ipsilateral asymmetries mainly reflect the anatomical wiring of the motor system, and only a fraction of the dominance of a given hemisphere leading to the preference of a given hand. Such a conclusion is also supported by the huge correlations between asymmetries of each hand movement showing that they were almost identical (R^2^ = 0.98 of right versus left hand asymmetries, supplementary Figure 1), as are the contralateral (R^2^=0.98) and ipsilateral (R^2^=0.96) variations.

Importantly, the support of handedness does not rely on a mirrored organization of the activity associated with preferred hand movement between RH and LH. Rather, it is associated with subtle activity modulations within the contralateral and ipsilateral cortices depending on the laterality status of the moving hand. As previously reported for the primary motor cortex (Tzourio-Mazoyer et al., 2015), handedness-modulated activity differences were actually seen in regions whose ipsilateral activity was related to difference between the preferred and non-preferred hand. But such a process was not limited to the primary motor cortex (rol3), since it was also true for SMA (cing4), although deactivation was stronger in LH, contrary to rol3. The present result, in line with previous studies establishing a link between ipsilateral deactivations and transcallosal inhibition from the dominant hemisphere (Hayashi et al., 2008; Tzourio-Mazoyer et al., 2015), shows that transcallosal inhibition varies locally with handedness, being more important in SMA (cing4) in LH. Such a spatial modulation within regions belonging to the same functional domain of motor control was also seen in somatosensory cortices. In INSP1 the deactivation triggered by the dominant hand was stronger than that of the non-dominant hand in RH than in LH, but the reverse was true in SMG1 and ROLop2. These findings demonstrate that subtle local variations of inhibition are coming from the hemisphere controlling the preferred hand, and that left-handedness cannot be reduced to a global decrease in inhibition, as we first proposed when examining the primary motor cortex alone (Tzourio-Mazoyer et al., 2015).

It is also important that the contralateral activations of Rolandic regions, INSP1 and subcortical areas were comparable in both groups during RFT and LFT, as that of the ipsilateral cerebellar activity, leading to opposite profiles when considering variations between preferred and non-preferred hand. Such an observation explains the difficulty in evidencing differences of task-related activations with handedness, and brings interesting information on the invariants for the right or the left hand movement, as it has been upper developed.

### Atlas of hand motor areas

In the line of the SENSAAS and WMCA atlases that are atlases we have proposed to the community on the networks dedicated to sentence and word-list processing, the 13 regions that have been selected and described in the present work are available as an atlas, the HAnd MOtor Area atlas (HAMOTA) at http://www.gin.cnrs.fr/en/tools/hamota.

## Data availability

Data have been submitted to Dryad (Mazoyer, Bernard et al. (2020), BIL&GIN FTT fMRI and handedness, Dryad, Dataset. DOI: 10.5061/dryad.cz8w9gj1z).

## Supplementary Material

**Supplementary Figure 1.**
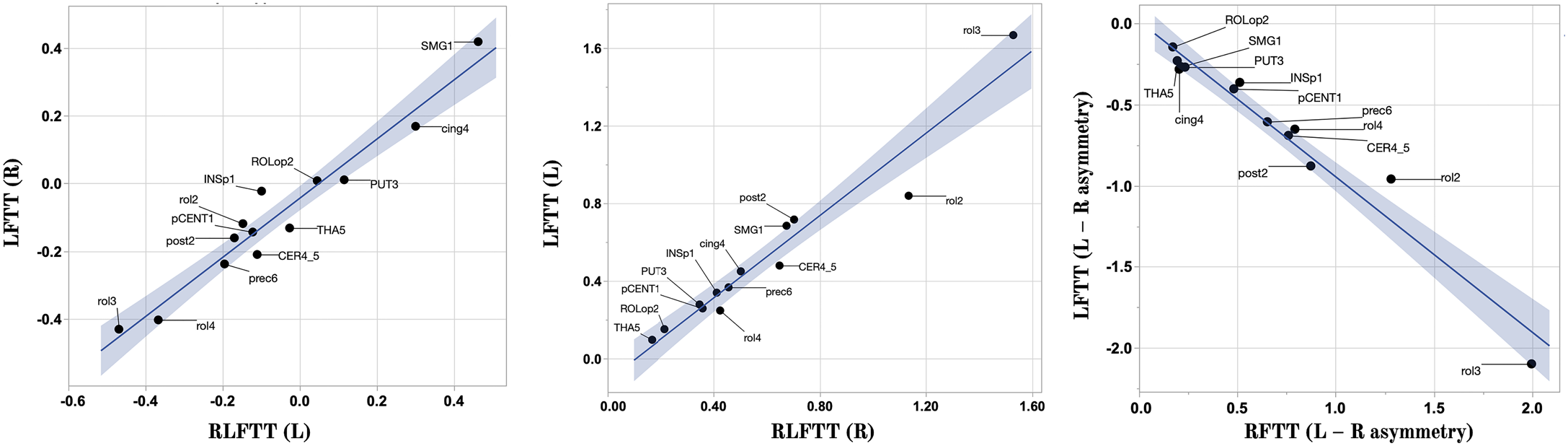
Regression of the contralateral mean activation and asymmetry values of each of the 13 hROIS in each task for the contralateral activation and asymmetries in the whole group. Activation strength across the 13 hROIs contralateral to the moving hand (ipsilateral for the cerebellar ROI) were very close during the RFT and the LFT (R = 0.94, left), as were ipsilateral activations (R = 0.94, middle), and asymmetries (R = 0.95, right).

